# Climatic limits for the present European distribution of hornbeam (*Carpinus betulus*), with special reference to Ukraine

**DOI:** 10.1101/506428

**Authors:** V.M. Tytar

## Abstract

In this study, we used an ecological niche modeling approach to detect the importance of diverse climatic parameters in controling the distribution of forest tree species, exemplified by the common hornbeam (*Carpinus betulus* L.), with special reference to Ukraine from where digitized data on the species has been scarce. In Ukraine populations of this tree species are found on the edge of its home range and are exposed to extreme climate conditions. The main objectives of the present study were to model the European-wide ecological niche of the common hornbeam and investigate primary climatic factors that control the potential distribution of this tree in Ukraine. Using an ecological niche modelling approach, we consider to have reliably modeled the European-wide bioclimatic niche of the common hornbeam for predicting the response of species’ geographical distribution to climate. Most contributing to the model were the mean monthly PET (potential evapotranspiration) of coldest quarter, continentality and annual PET. In terms of the “Most Limiting Factor”, in Ukraine continentality is crucial throughout the majority of the country.

## Introduction

Forests, which today cover 30.8 percent of the world’s land surface (MacDicken et al., 2015), are being rapidly and directly transformed in many areas by ongoing climatic changes and impacts of expanding human populations and economies. Forest decline commonly involves multiple, interacting factors, ranging from drought to insect pests and diseases, often making the determination of a single cause unrealistic. However, abiotic stress factors are commonly responsible, with climate stresses thought to be a primary triggering factor (Desprez-Loustau et al., 2006; Raffa et al., 2008).

It is well known that the macrodistribution of plants is primarily controlled by climate (Woodward, 1987; Prentice et al., 1992). Vegetations have long been classified on the basis of climate, which results in several classification systems (e.g. Holdridge, 1947; Thornthwaite, 1948; Kira, 1976). Of various climatic factors, heat sum, water availability, and winter coldness have been suggested as primary limitations for plant distribution over the large scale (Woodward, Rochefort, 1991).

Trees are foundational elements of forest ecosystems (Ellison et al., 2005), having important influences on the resource environment and ecological function, and understanding the relative roles of climate in controlling tree species distribution is a basis for predicting responses to potential climatic changes (Loehle, LeBlanc, 1996; Austin, 2002).

Because the present distributions of most tree species are strongly related to climate, long ago simple isotherm methods and/or comparisons of climatic maps with geographic ranges have been used to assess distributional and limiting factors in relation to climate (i.e. Hutchinson, 1918; Koppen, 1936; Huntley et al., 1995; Sykes et al., 1996; Cimhob, Дiдyx, 2001), however using these methods alone may be insufficient to explore the nuances of climatic controls on range limits of species. Today species distribution models (SDMs) are the prime tools used to assess the spatial distribution of potentially suitable habitat for species, and to hypothesise how suitability is affected by the environment (Guisan, Thuiller, 2005). With the development of computer hardware and Internet speed, the current availability of species and climate information sharing systems has greatly enhanced the field of SDM. These tools generally correlate species’ occurrence patterns with environmental variables, which are frequently selected from an array of ‘bioclimatic’ indices. In this study, we used this comprehensive approach to detect the importance of diverse climatic parameters in controling the distribution of forest tree species, exemplified by the common hornbeam (*Carpinus betulus* L.), a notable component of many European forest ecosystems, with special reference to Ukraine from where digitized data on the species has been scarce.

Common hornbeam is a cool summergreen nemoral mesophyte deciduous (Laurent et al., 2004), usually considered late-successional tree, that frequently forms part of several types of mixed broad-leaf communities in European temperate summergreen forests. Forests of such kind are currently confined between ca. 55–60° and 40°N, distributed in the Boreal, Atlantic, Alpine, Mediterranean, Continental and Pannonian floristic regions (European Commission 2013). They are predominant in western and central Europe but also extend through Eastern Europe to the Ural Mountains (Hendrick, 2001). This large area of distribution broadly agrees with the main range of the common hornbeam, except for its absence in most of Iberia (Sikkema et al., 2016) and in areas stretching from Eastern Ukraine towards the Urals (Bopoбйob ta ih., 2008; Удpa, 2010). Currently, *C*. *betulus* is only very locally distributed in Eastern Ukraine, but it has been assumed that there were several areas suitable for the persistence of *C*. *betulus* in the past (Фeдopoba, 1951). Major environmental changes, coupled with human impact (Удpa, 1988; Pokorný, 2005; Tonkov et al., 2013), are explanations of the progressive decline of *C*. *betulus* in Europe during the Late Holocene, and suggesting that modern hornbeam populations occurring in Eastern Ukraine could be considered a relict of a larger distribution in the past. In Ukraine populations of common hornbeam found on the edge of its home range in forest-steppe ecotones are exposed to an extreme climate with water shortages during the growing season and low temperatures in winter. Usually forests in these ecotones are highly fragmented and, in addition to climate, are heavily subjected to land use.

The main objectives of the present study were to model the European-wide ecological niche of the common hornbeam and investigate primary climatic factors that control the potential distribution of this tree in Ukraine.

## 2. Material and Methods

### 2.1 Species Occurrence Data

Georeferenced occurrence data were collected from the Global Biodiversity Information Facility (*Carpinus betulus* L. in GBIF Secretariat, 2017) and the EU-Forest, a high-resolution tree occurrence dataset for Europe (Mauri et al., 2017). Unfortunately, these databases hold hardly any georeferenced records of the common hornbeam in Ukraine, therefore a search was undertaken to complement the information with point data from Ukraine using available internet resources, particularly of the State Forest Resources Agency of Ukraine (http://dklg.kmu.gov.ua), and published scientific literature containing explicit location data regarding the species (i.e. Bopoбйob ta ih., 2008; Удpa, 2010; Maлa, 2012 etc.). This produced 124 georeferenced records for mainland Ukraine. In total 13,347 collected records were considered.

Not surprisingly, these occurrence points varied in spatial density due to variable sampling intensity over geography. As a result, and to avoid overemphasizing heavily on sampled area, we selected points for model calibration using a two-step subsampling regime to reduce sampling bias and spatial autocorrelation, which would produce models of lower rather than higher quality (Beck et al., 2013). In the first step we collated records with a moderate coarse geographic resolution of 5 arc-min (0.08333°) using ENMTools (Warren et al., 2010). The main reason for this was that there could have been a sampling bias or error at a fine resolution (particularly in the GBIF database). Secondly, we used the OccurrenceThinner software (version 1.04) (Verbruggen et al., 2013), which reduces geographical bias in datasets by a probability-based procedure defined by a kernel density grid created in SAGA GIS software (Conrad et al., 2015). This considerably reduced the original dataset to 6,658 records.

The algorithms used in this paper, Maxent (Phillips et al., 2006) and BIOCLIM (Nix, 1986), have proven good performance and accuracy for such studies (Elith et al., 2011).

Maxent (Version 3.3.3k) is a machine learning algorithm (Phillips et al., 2006). The main advantage of applying Maxent to the modeling of geographical species distributions in comparison with other methods is that it only needs presence data, besides the environmental layers. Pseudo-absence points (used instead of true absences) were randomly generated within a convex hull encompassing hornbeam presence points.

Maxent provides output data in various formats. The logistic format (values range from 0 to 1) is recommended, because it allows for an easier and potentially more accurate interpretation of habitat suitability compared to the other approaches (Baldwin, 2009). Maps of habitat suitability were imported into SAGA GIS and DIVA GIS (Hijmans et al., 2001).

Maxent modeling can determine the importance of environmental variables. In one option the contribution for each variable can be determined by randomly permuting the values of that variable and measuring the resulting model performance. Better performance means that the model depends heavily on that variable (for explicitness values are normalized to give percentages). The second option uses a jackknife test and the regularized gain change during each iteration of the training algorithm. The environmental variable with the highest gain is considered to have the most useful information by itself, whereas the variable causing the largest decrease in the model’s gain contains the most information not found in the other environmental variables. Both of these options are used to determine the importance of the environmental variables and to observe whether or not the predictive distribution responded to each variable as expected.

Maxent also allows the construction of response curves to illustrate the effect of selected variables on habitat suitability (consequently, on the probability of occurrence and giving an idea of where for each variable, under the constraints and conditions of the modelling situation, the focal species has it’s optimum). These response curves consist of the specific environmental variable as the x-axis and, on the y-axis, the predicted probability of suitable conditions as defined by the logistic output. Upward trends for variables indicate a positive relationship; downward movements represent a negative relationship; and the magnitude of these movements indicates the strength of the relationship (Baldwin, 2009).

An important part of determining the ability of niche models to predict the distribution of a species is having a measure of fit. The performance of the Maxent model is usually evaluated by the threshold-independent receiver operating characteristic (ROC) approach (calculating the area under the ROC curve (AUC) as a measure of prediction success). The ROC curve is a graphical method that represents the relationship between the false-positive fraction (one minus the specificity) and the sensitivity for a range of thresholds. It has a range of 0–1, with a value greater than 0.5 indicating a better-than-random performance event (Fielding, Bell, 1997). A rough classification guide is the traditional academic point system (Swets, 1988): poor (0.5–0.6), fair (0.6–0.7), good (0.7–0.8), very good (0.8–0.9) and excellent (0.9–1.0). A 10-fold cross validation was used to evaluate the fit goodness of the Maxent model in simulating the current potential suitable habitat with occurrence data and variables for the current environment. This method randomly partitions the original occurrence records into 10 subsets of equal size. Of the 10 subsets, one subset was used as testing data, while the remaining nine subsets are used as training data.

Maxent uses “regularization” (the beta parameter) to avoid over-fitting data. Higher values of beta increase the “smoothness” of species’ responses to the environment. The default value of beta in Maxent is 1, however we applied values of 1, 2, 3 and 4. The final models have been selected, which incorporate a lesser number of correlated variables, and an optimum value of beta (see below).

The “Most Limiting Factor” output, available in BIOCLIM (implemented in DIVA GIS), was also applied to detect which variable in Ukraine is the most critical for the inclusion of the grid cell within the resulting environmental envelope. The “Most Limiting Factor” model outputs the variable with the lowest or highest score in each grid cell, for which there is a prediction assigning them a certain colour on the prediction map. For the cells that fall within the 0–100 percentile, the variable with the lowest percentile score is mapped.

### 2.2. Environmetal variables

According to previous studies, there are many variables (including hydrological-thermal) that are thought to be relevant to species’ ecology and geographic distribution. Nevertheless, the selection of biologically meaningful environmental variables that determine relative habitat suitability is a crucial aspect of the modeling pipeline. In this study we have used a recently reconsidered (in terms of biological significance) set of 16 climatic and two topographic variables (the ENVIREM dataset), many of which are likely to have direct relevance to ecological or physiological processes determining species distributions (Title, Bemmels, 2018).

The most important set of variables assessed in performance of models was identified using the *Maxent Variable Selection* package in R developed by (Jueterbock et al., 2016). The program first tests for Pearson correlation, then through an iterative process tests different regularization values, correlation and contribution of variables to determine the best possible set of variables by sample size corrected Akaike information criteria, AICc (Akaike, 1974). This procedure resulted in the selection of 7 climatic variables included for further analyses (Table 1).

**Table 1.** Environmental variables used in this study and their permutation contribution to the model.

| Variable abbreviation | Brief description | Units | Contribution (%) |
| --- | --- | --- | --- |
| aridityIndexThornthwaite | Thornthwaite aridity index: Index of the degree of water deficit below water need. | - | 5.1 |
| continentality | Average temperature of warmest month minus the average temperature of coldest month. | °C | 26.7 |
| growingDegDays5 | Sum of mean monthly temperature for months with mean temperature greater than 5 °C multiplied by number of days. | - | 8.4 |
| maxTempColdestMonth | Maximum temperature of the coldest month. | °C*10 | 8.1 |
| minTempWarmestMonth | Minimum temperature of the warmest month. | °C*10 | 1.8 |
| annPET | Annual potential evapotranspiration: a measure of the ability of the atmosphere to remove water through evapotranspiration processes, given unlimited moisture | mm/year | 21.1 |
| PETColdestQuarter | Mean monthly PET (potential evapotranspiration) of coldest quarter. | mm/month | 28.7 |

## 3. Results

The best model with fewer predictors (see above) for the European-wide ecological niche of common hornbeam was obtained when the beta parameter equaled the default value of 1.0 (minimum AICc=139318.3). The Maxent model showed a reliable (or “good”) prediction with an AUC of 0.726 (SD=0.007), greater than the 0.5 of a random model.

In terms of permutation importance, “PETColdestQuarter” contributed most to the model (28.7%), followed by, “continentality” (26.7%), and “annPET” (21.1%) (Table 1). These three variables collectively contributed 76.5% of the variance in the bioclimatic model simulating the geographical distribution of *C*. *betulus* in Europe.

The leading role of the first two was accordingly supported by the jackknife tests of variable importance: the environmental variable with highest gain when used in isolation is “PETColdestQuarter”, which therefore appears to have the most useful information by itself, whereas the environmental variable that decreases the gain the most when it is omitted is “continentality”, which therefore appears to have the most information that isn’t present in any of the other variables used for building the model. The remaining four bioclimatic factors were less important in determining the geographical distribution of *C*. *betulus* (collectively, they explained 23.4% of the variance, 1.8%–8.4% for each factor). The response curves of first three most important climatic factors (PETColdestQuarter, continentality and annPET) are presented in Figure 1, showing a general hump-shaped relationship between the habitat suitability values and values of the corresponding climatic factors. Using a 50% habitat suitability threshold (Waltari, Guralnick, 2009), suitable conditions for the distribution of the common hornbeam in Europe (in terms of the ENVIREM dataset) are PETColdestQuarter values ranging between 12.8-18.06 mm/month, continentality (15.07-18.86°C), and annPET (706.6-902.3 mm/year).

**Fig 1.**
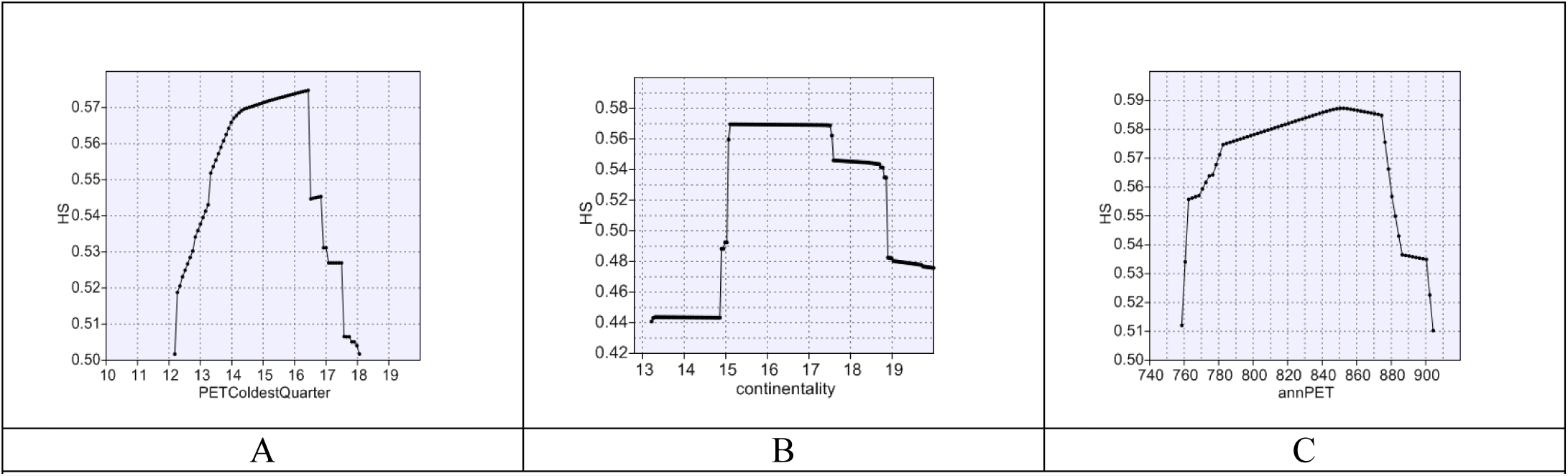
The response curve of bioclimatic habitat suitability (HS) for three dominant climatic factors based on the *Maxent* model created using only the corresponding variable; (A) PETColdestQuarter (mm/month), (B) continentality (°C), (C) annPET (mm/year).

In terms of the “Most Limiting Factor” output (Fig.2), in Ukraine the most critical for the inclusion of a grid cell within the resulting environmental envelope was the “continentality” predictor. For the majority of the country (around 78%), including northerly portions of the Crimea, Transcarpathia and NW Volhyn’, this is a crucial climatic factor limiting the distribution of common hornbeam. In a considerably smaller portion of the country (9.8%), namely in western Ukraine and southern parts of the Crimea, suitable conditions for the species are limited by either “PETColdestQuarter” or “annPET”, apparently depending on the topography of the area. Further to the east conditions are predicted to be outside of the European bioclimatic envelope of *C*. *betulus* and can be considered unsuitable for the species; this area occupies around 12.2% of the country.

**Fig. 2.**
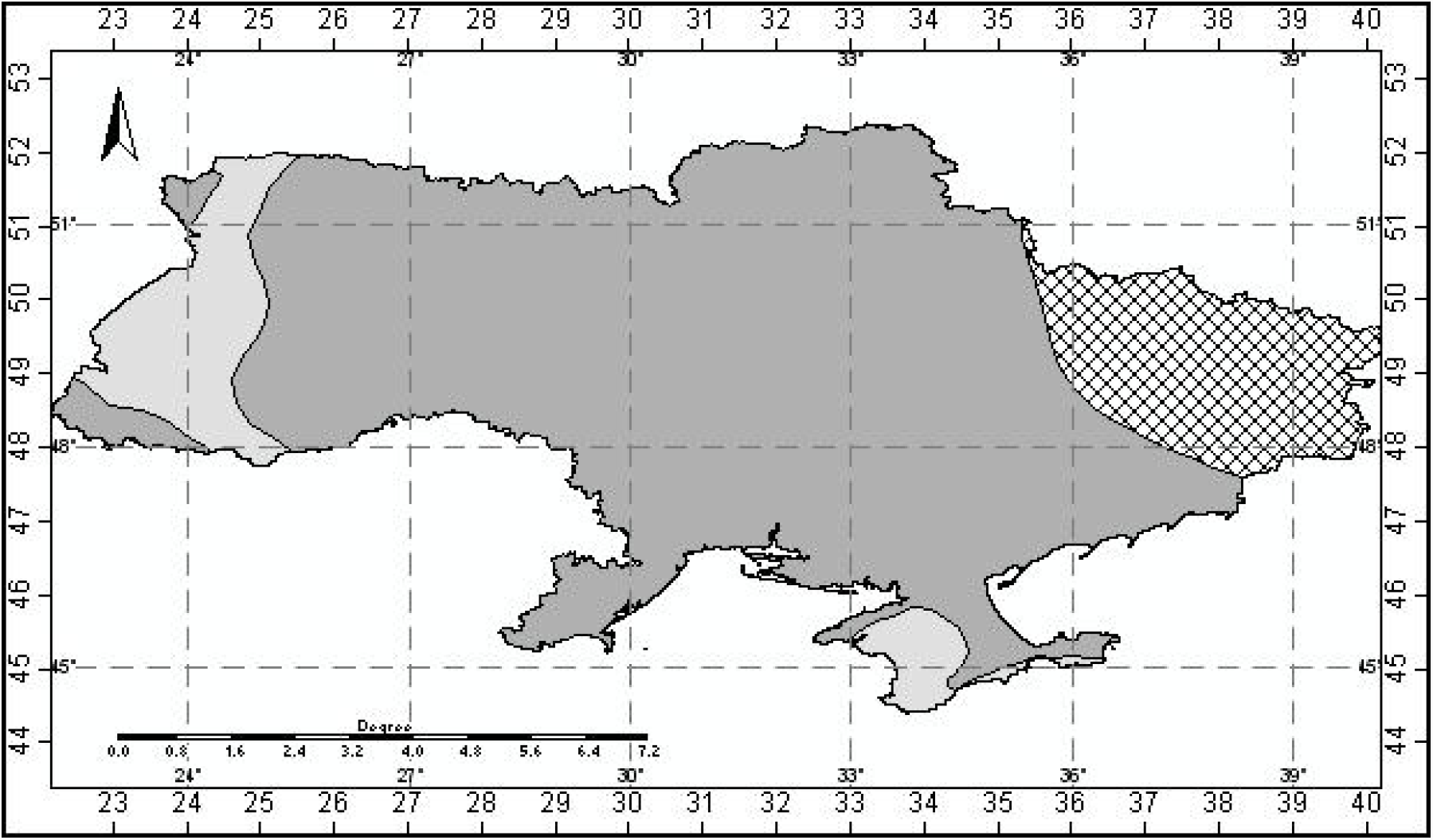
The “Most Limiting Factor” model for Ukraine. Gray shading indicates the geography of the suitability values within the bioclimatic envelope of *C*. *betulus*. Cross diagonal filling shows the area where conditions are outside the bioclimatic envelope of the species. The darker shade indicates areas with suitability limited by “continentality” while the lighter shade denotes areas where “PETColdestQuarter” and “annPET” are limiting factors.

## 4. Discussion

Climate is considered to be the most important environmental factor influencing the distribution of species and vegetation at a regional and global scale. Our research at the regional scale (Ukraine) has concentrated on investigating the climatically suitable habitat, significant climatic factors, climatic thresholds (niche) and climatic response curves of the common hornbeam using a European-wide approach. The “good” prediction of the resulting *Maxent* model (with an AUC of 0.726) may seem insufficient, however it could be that species (such as *C*. *betulus*) with larger areas of occupancy, e.g. larger range size, and those found over a greater range of environmental conditions, yield models with lower accuracy than those with smaller ranges (Franklin et al., 2009; Gray, Hamann, 2013). Besides this, climate variables are unlikely to be the only relevant predictors of habitat suitability (Austin, Van Niel, 2011), as plant survival and reproduction also depends on light, nutrients, water, soil type and CO_2_, as well as disturbances and biotic interactions (Hageer et al., 2017). Including these variables could have improved model performance, but here we explicity focus on climate. Nevertheless, using only climate variables the modelling exercise gave accurate projections of common hornbeam distribution beyond the convex hull (used for calibrating the model; see above). For instance, the *Maxent* model simulated potential habitat for the species in the Crimea, where hornbeam is found in isolation (Дидyx, 1992; Cordova, 2016).

In terms of variable importance, around 50% of the variation in the distribution of the study species is due to measures of evapotranspiration (Table 1). Evapotranspiration has long been regarded as an index to represent the available environmental energies and ecosystem productivity by bioclimatologists (Sun et al., 2016) and has been used to explain large regional variations in plant and animal species’ richness and biodiversity. For example, the variability in species richness in vertebrate classes could be statistically explained by a monotonically increasing function of a single variable, potential evapotranspiration (PET) (Currie, 1991). In contrast, regional tree richness was more closely related to actual evapotranspiration, AET (Hawkins et al., 2003). To relate these findings to our study, we assume that geographical variations in habitat suitability for a separate species (as *C*. *betulus*) and such variations in species’ richness must have at least some common drivers as far as both are dependent on ecologically important aspects of climate linked to energy supply (O’Brien, 2006), measured by PET (Fisher et al., 2011). However, despite this importance, a quantitative synthesis analysing the contribution of over 400 distinct environmental variables to 2040 *Maxent* species’ distribution models PET is poorly represented: summer PET is accounted for in 34 articles whereas winter PET only in three (Bradie, Leung, 2017). Nevertheless, winter PET in our case (represented by “PETColdestQuarter”) turned out to be a highly influential predictor of the European distribution of the common hornbeam.

PET is minimal for whole study region in winter and the main explanation for that is the insufficiency of solar energy during the winter. Yet there seems to be a crucial demand by the common hornbeam in the colder quarter of the year when trees are considered to be dormant. Dormancy is usually stated as a period in which visible growth is not obviously apparent (Saure, 1985), but cell division and differentiation, tissue growth continue to occur at low temperatures in a slow and steady manner, and there is also active synthesis of RNA and protein during dormancy (Bagni et al., 1977). The corresponding response curve (Fig.1A) may indicate the need for a sufficient energy supply to support these processes. Consequently, areas with a better such supply are of better habitat suitability, which reaches a peak at 0.57 by “PETColdestQuarter” value of 16.43 mm/month. Further, however, the response curve shows a sudden drop. Most likely this indicates that a surplus of ambient energy may have the potential to break dormancy and considerably reduce habitat suitability by causing premature growth onset and heavy damage during subsequent periods of frost, a possible scenario that may occur under climate change (Hänninen, 1991).

Likewise the “PETColdestQuarter” predictor, annual potential evapotranspiration (“annPET”), an index representing accumulated energy availability throughout the year (Thornthwaite 1948), substantially contributes to the *Maxent* model, and is strongly correlated to potential evapotranspiration occurring in the warmest quarter (summer). Excessive values of “annPET” apparently lead to a deficit of moisture in the soil, therefore after a value of 876.36 mm/year there is a sharp decline of the corresponding response curve (Fig.1C) indicating worsening habitat suitability for hornbeam in areas where “annPET” values exceed this threshold, and this could be due to the summer water deficit which is damaging for hornbeam populations (Sykes et al., 1996).

In contrast to “PETColdestQuarter” and “annPET”, “continentality” causes the largest decrease in the model’s gain therefore includes information not found in the first two. “Continentality”, as a distinguishable dimension of the climatic niche of the species, features the seasonal amplitude in ambient temperature, where large differences between low temperatures (frost) in the winter, and high temperatures (heat) in June, are suggested to be limiting factors of tree growth (Clark et al., 2014; Augustaitis et al., 2015). According to Sykes et al. (1996), common hornbeam can tolerate winter mean temperatures as low as –8°C. Using GIS to extract climate data from the CliMond dataset (Kriticos et al., 2012), we arrived with approximately the same figure (–8.65°C) for mean temperatures of the coldest quarter, but also found the first quartile to be close to zero (precisely –0.8 °C), meaning only 25% of the point data are from places where the mean temperatures of the coldest quarter are below this level. Interestingly, values at the lower extremes of “continentality” (in fact, increased “oceanicity”) are just as restraining as those of the upper (Fig.1 B), indicating that there is perhaps some “degree of continentality” essential for the well-being of the species. This notion has been much earlier expressed by D.D. Lavrinenko, a Soviet botanist, who wrote “… each of the climatic forest zones (climatopes) should be described by the definite amount of warmth…received by the plants and by the definite degree of continentality of climate…” (Lavrinenko, 1972).

Because the distribution of any species depends on variables related to climate, it is likely that the species could rapidly respond to climatic change (Luoto et al. 2006). In this respect the common hornbeam is no exception, as exemplified by the history of the species in Eastern Europe. Limiting factors and thresholds (particularly of PET indices) are are bound to shift together with global climate change and bring in changes to the pattern of the common hornbeam distribution, especially at its edges.

## 5. Conclusions

Species distribution models are powerful tools to assess the potential geographical distributions of target species, relying on the correlation between species existence and corresponding environmental variables. Using the ENVIREM dataset (Title, Bemmels, 2018), presence-only point data (supplemented by data from Ukraine), and by applying a maximum entropy (*Maxent*) model approach (Phillips et al., 2006), we consider to have reliably modeled the European-wide bioclimatic niche of the common hornbeam for predicting the response of species’ geographical distribution to climate. Most contributing to the model were the mean monthly PET (potential evapotranspiration) of coldest quarter, continentality and annual PET. In terms of the “Most Limiting Factor”, in Ukraine continentality is crucial throughout the majority of the country.

